# KRAB Zinc Finger protein Znf684 interacts with Nxf1 to regulate mRNA export

**DOI:** 10.1101/2021.09.29.462476

**Authors:** Alexandra Nitoiu, Syed Nabeel-Shah, Shaghayegh Farhangmehr, Shuye Pu, Ulrich Braunschweig, Benjamin J. Blencowe, Jack F. Greenblatt

## Abstract

Cys2His2 (C2H2) type zinc finger (ZnF) proteins constitute a large class of proteins that are generally considered to be DNA-binding transcription factors. Using affinity purification followed by mass spectrometry, as well as reciprocal co-immunoprecipitation experiments, we determined that the C2H2-ZnF protein Znf684 interacts physically with several proteins involved in mRNA export, including Nxf1 and Alyref. We utilized individual nucleotide resolution cross-linking immunoprecipitation followed by high throughput sequencing (iCLIP-seq) experiments to show that Znf684 binds directly to specific mRNAs *in vivo* and has an RNA-binding profile similar to those of Nxf1 and Alyref, suggesting a role in mRNA export regulation. Immunofluorescence microscopy (IF) experiments revealed that Znf684 localizes to both the nucleus and cytoplasm. Using cellular fractionation experiments, we demonstrate that overexpression of Znf684 negatively impacts the export of SMAD3 and other target mRNAs. Taken together, our results suggest that Znf684 regulates the export of a subset of transcripts through physical interactions with Nxf1 and specific target mRNAs.

## Introduction

Gene expression regulation is critical in maintaining proper cell function, and its misregulation has been implicated in the development of various pathologies and diseases (1). Recent studies indicate that selective export of mRNAs is one of the major posttranscriptional regulatory mechanisms (2–5). For mRNA to be successfully exported from the nucleus to the cytoplasm and translated by ribosomes, it needs to go through several steps. First, it must be properly processed, undergoing capping, splicing, and 3’ end processing (6). Second, it needs to associate with proteins to become a messenger ribonucleoprotein (mRNP) (7, 8). Lastly, it must dock at a nuclear pore complex (NPC) and travel through it to reach the cytoplasm (6). Once the mRNA enters the cytoplasm, the mRNA export factors dissociate from it, ensuring that the mRNA does not return to the nucleus (7).

There are several different export pathways which mRNAs may use to reach the cytoplasm, and the two most common ones rely on the proteins Nxf1 and CRM1 (6). One of the proteins that is required for mRNPs to be transported by NPCs is the Tap-p15 heterodimer, also known as Nxf1-Nxt1 in animals (7). Although Nxf1-Nxt1 binds non-specifically to RNA, specific mRNA-binding proteins are thought to function as adaptors (7). The transcription-export complex (TREX), which includes the RNA-binding proteins Alyref and Chtop (2), interacts with Nxf1-Nxt1 heterodimers, functioning as an adaptor to ensure that these heterodimer bind specifically to mRNA (7). TREX is recruited to the 5’ end of mRNA during capping and splicing, but before 3’ end processing (2). The cap binding complex (CBC) transiently binds to Alyref, and then Alyref transfers to sites located beside exon junction complexes (EJC) throughout the mRNA during co-transcriptional splicing (2). On the other hand, Chtop preferentially binds to 3’ untranslated regions (UTRs) and regulates alternative polyadenylation (2). TREX then recruits Nxf1 to mRNAs after they have been fully processed (6). The Nxf1 nuclear export pathway has been shown to have plasticity, since it can be used to transport specific mRNA cargoes, and Nxf1 can interact with different adaptor proteins or with the mRNA directly to achieve this (6). Although various studies have shown that Nxf1 is necessary for RNA nuclear export, Alyref is not, as other factors can also function as adaptors between the mRNA and Nxf1 (9). Since recent studies suggest that specific biological pathways can be regulated by selective mRNA export (5), it is conceivable that there are additional Nxf1 adaptor proteins which have not been identified.

Although nuclear mRNA export is important for eukaryotic cell function, it can also be vital for viral infection and proliferation (7, 10). For example, it has been demonstrated that severe acute respiratory syndrome coronavirus 2 (SARS-CoV-2) disrupts the normal nuclear export pathway in several ways (10, 11), such as by interacting with the Nxf1-Nxt1 heterodimer to prevent mRNA export from the nucleus (11). Additionally, various studies have shown that several factors involved in mRNA nuclear export, including Nxf1 and Alyref (12), are dysregulated in cancer as well as in certain genetic diseases (9), indicating the broad relevance of research on this topic.

Cys2His2 (C2H2) type zinc finger (ZnF) proteins are a large class of proteins (>700 in humans) (13) that proliferated in primates due to gene duplication (14). Krüppel-associated box (KRAB)-containing zinc finger proteins (KZFPs) comprise roughly half (~350) of the human C2H2-ZnFs. KZFPs are capable of recruiting heterochromatin-inducing KAP1 (TRIM28) and are thought to function as repressors of transposable elements (TEs) during early embryogenesis (15). Although C2H2-ZnFs are generally considered to be DNA-binding transcription factors (13, 16–18), some of these proteins (e.g. CTCF, YY1 and GTF3a) have also been shown to function as RNA-binding proteins (RBPs) (19, 20). This suggests that C2H2-ZnFs might have functions that are not yet known. Here, we show that a previously uncharacterized KZFP, Znf684, physically interacts with several proteins involved in mRNA export, including Nxf1 and Alyref. Our *in vivo* RNA-binding data indicates that Znf684 binds directly to specific mRNAs and has a binding profile similar to those of Nxf1 and Alyref. We also found that overexpression of Znf684 plays an inhibitory role in the mRNA export of certain target transcripts including SMAD3, VEGFA, DDX3X and AKT1. Together, our results suggest that Znf684 functions in mRNA export regulation.

## Results

### Znf684 is conserved across vertebrates

We initiated our investigation by studying the sequences of Znf684 and its orthologues in other species. We found that Znf684 has 8 Zinc finger domains, in addition to a KRAB domain at its N-terminus. Znf684 is very highly conserved in primates, such as humans, chimpanzees, and gorillas, and quite well conserved in some other vertebrates, such as horses and dogs, with protein sequence identity > 65% (Figure 1A). We observed that certain species, such as dogs, have lost the KRAB domain, resulting in a decrease in the protein length by approximately 30 residues (Figure 1A). However, the number and placement of the zinc finger domains remains evolutionarily conserved. This observation suggests that Znf684 might be functionally divergent in species lacking the KRAB domain, perhaps not functioning to repress gene expression. Of note, certain species, such as mice and ray-finned fish, appear to have lost the *ZNF684* gene entirely.

**Figure 1:**
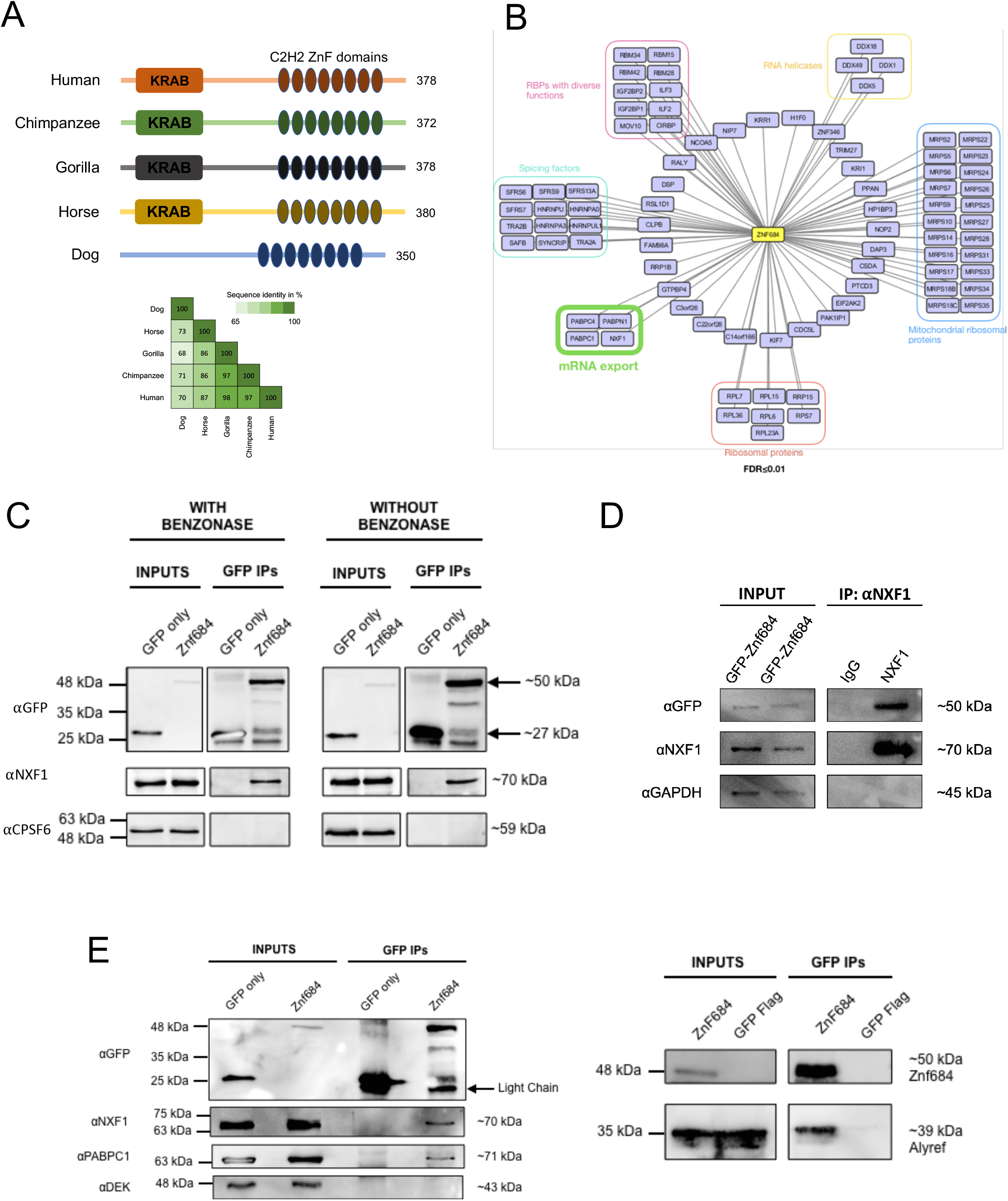
Znf684 interacts with mRNA export factor Nxf1. **A**: Comparative domain analysis of human Znf684 protein against primate, horse, and dog orthologues. Overall sequence identity among the orthologues is shown in the panel below. **B:** Network representation of high confidence (FDR ≤ 0.01) Znf684 protein-protein interactions. Bait node is shown in yellow and the copurifying partners are grouped according to their known roles. The co-purifying partners associated with mRNA export are highlighted in green. **C:** Co-IPs were performed using whole cell lysates prepared from HEK293 cells expressing either free GFP or GFP-Znf684. IPs were performed using anti-GFP antibody. The blots were probed with the indicated antibodies, with CPSF6 acting as loading control. **D:** Reciprocal co-IPs were performed using anti-Nxf1 antibody. The blots were probed with the indicated antibodies. **E:** Co-IP analyses using whole cell lysates prepared from HEK293 cells expressing either free GFP or GFP-Znf684. The blots were probed with the indicated antibodies.

We next examined the Znf684 expression profile across various tissues and cell lines using publicly available RNA sequencing (RNA-seq) data. In general, Znf684 appears to be weakly expressed across the tissues and cell lines we examined (Supplementary figure 1A). However, Znf684 is significantly upregulated during embryonic development at stages corresponding to 8C and morula (Supplementary figure 1B). These observations suggest that Znf684 expression is tightly regulated across tissues and during embryonic development.

### Znf684 interacts physically with mRNA export factors

To begin characterizing Znf684, we analyzed our previously published affinity purification followed by mass spectrometry (AP-MS) data generated in human embryonic kidney (HEK293) cells inducibly expressing GFP-tagged variant of Znf684 (21). We scored Znf684 AP-MS data against numerous control purifications using Significance Analysis of INTeractome express (SAINTexpress) (22). Application of SAINTexpress indicated that Znf684 binds to mRNA nuclear export factors such as Nxf1 and Pabpn1, as well as to mitochondrial ribosomes and other proteins (False Discovery Rate [FDR] <0.01; Figure 1B, Supplementary Table S1). This suggested that Znf684 might have roles in mRNA export regulation and mitochondrial translation.

To validate the interaction with mRNA export factors, we performed coimmunoprecipitation (co-IPs) experiments using whole cell extracts prepared from cells expressing either free GFP or GFP–Znf684 and probed the resulting Western blots with antibody against Nxf1. Cell lysates were treated with a promiscuous nuclease (Benzonase) to reduce any indirect RNA (or DNA)-mediated interactions, ensuring that only physical protein-protein interactions were captured in co-IPs. Our results show that Nxf1 is present only in the GFP-Znf684 immunoprecipitated samples and not in the free GFP ones (Figure 1C). To further validate these findings, we performed reciprocal Co-IP experiments and observed that endogenous Nxf1 was able to pull down GFP-Znf684 (Figure 1D).

We next asked whether or not Nxf1 could also interact with Alyref and PABPC1, which have also been shown to function in the mRNA export pathway (2, 23). As shown in figure 1E, Znf684 successfully pulled down both Alyref and PABPC1. These results confirmed the mass spectrometry data and showed that Znf684 interacts physically with mRNA export-related proteins. Given that Znf684 interacts with both Nxf1 and components of the TREX complex, it became likely that Znf684 plays a role in mRNA export regulation.

### Znf684 is present in both the nucleus and mitochondria

To begin understanding the mechanism of Znf684’s function in mRNA export, we studied Znf684’s subcellular localization. Using publicly available immunofluorescence (IF) data from the Human Protein Atlas (www.proteinatlas.org), we observed that endogenous Znf684 localizes to both the nucleus and cytoplasm in several human cancer cell lines, including Cervical cancer (SiHa), Bone Osteosarcoma Epithelial (U2OS), and Rhabdomyosarcoma (RH30) cells (Supplementary Figure S2). We also performed IFs in HEK293 cells overexpressing either GFP-Znf684 or free GFP. Our results showed that Znf684 localizes to both the nucleus and cytoplasm, consistent with its localization in human cancer cells (Figure 2A). Since the cytoplasmic signal appeared as distinct puncta, bearing resemblance to mitochondria, and Znf684 interacts with many mitochondrial ribosomal proteins, we examined whether Znf684 localizes to mitochondria by co-staining cells with the mitochondrial marker Mitotracker. We found that Znf684 does localize to mitochondria, in addition to its presence in the nucleus (Figure 2B).

**Figure 2:**
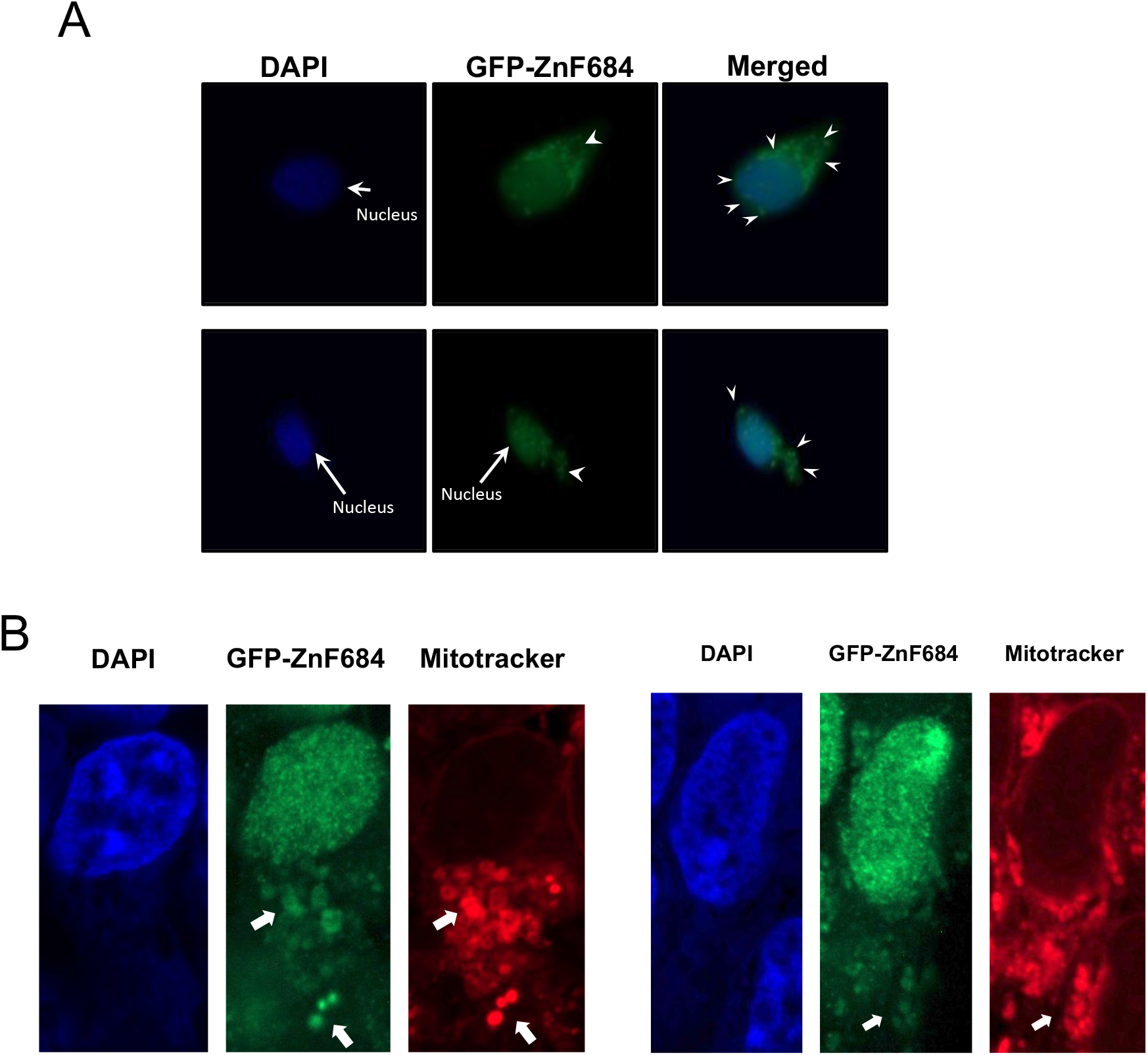
Znf684 localizes to the nucleus and mitochondria. **A**: IF analysis of GFP-Znf684 in HEK293 cells. DAPI was used for nuclear staining. White arrows point towards the nucleus. White arrowheads point towards puncti and protrusions of GFP staining that appear beyond the nucleus. **B:** IF analysis of GFP-Znf684 in HEK293 cells. DAPI was used for nuclear staining and Mitotracker was used for mitochondrial staining. White arrows point towards mitochondria.

### Znf684 binds directly to RNA

To investigate whether Znf684 binds directly to RNA, we performed crosslinking and immunoprecipitation (CLIP), followed by gel electrophoresis and autoradiography. Our data shows that GFP-Znf684 cross-linked robustly to RNA in UV-irradiated cells, while GFP alone did not crosslink and thus did not produce any observable radioactive signal (Figure 3A). Since the strength of the radioactive signal decreased in response to over-digestion with RNase I, we concluded that Znf684 binds directly to RNA (Figure 3A).

**Figure 3:**
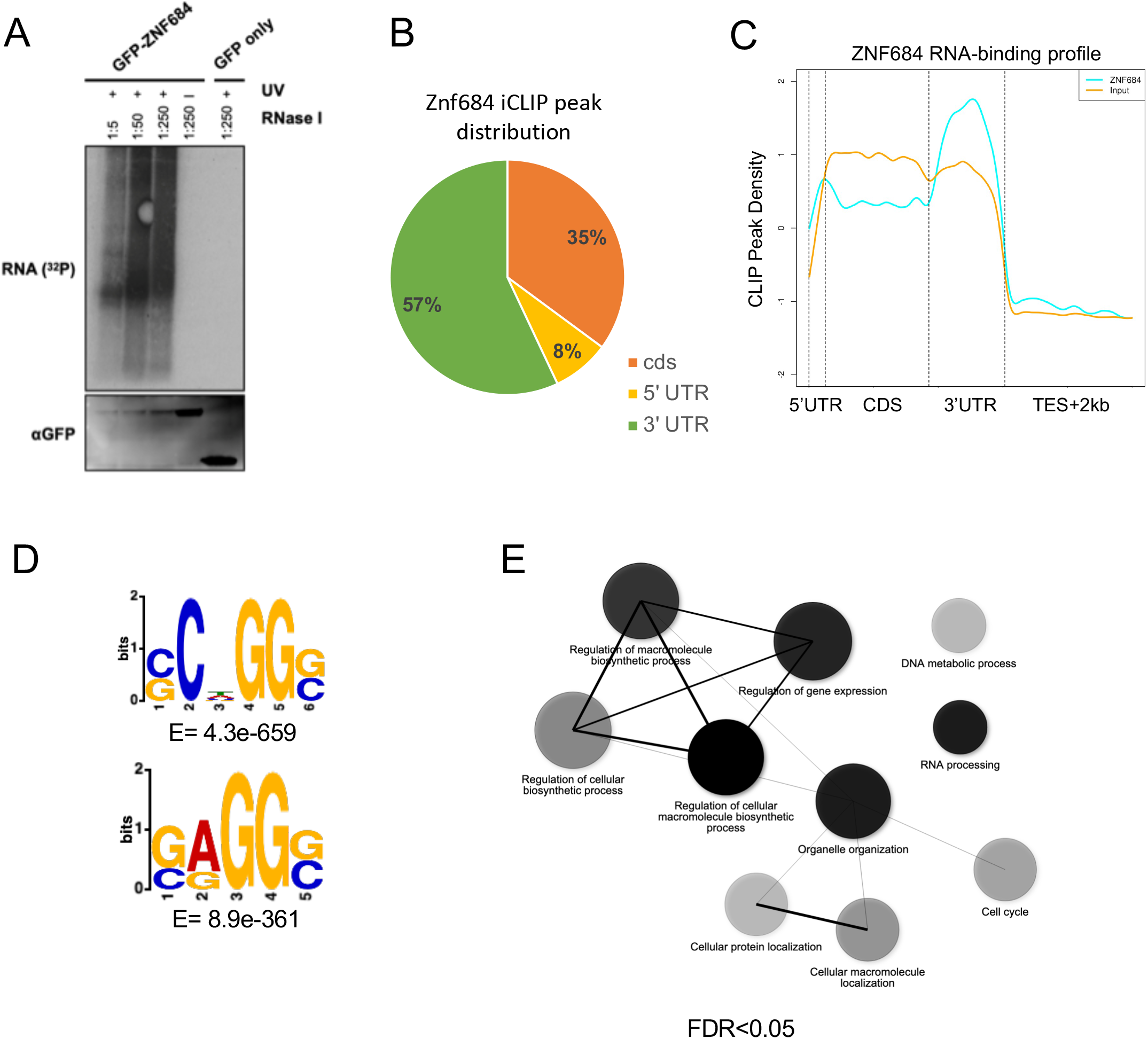
Znf684 directly binds RNA. **A:** The top panel shows autoradiographs of immunopurified ^32^P-labeled Znf684-RNA complexes after partial RNase I digestion. HEK293 cells were UV-crosslinked and GFP-Znf684 was immunoprecipitated using anti-GFP antibody. After radiolabeling the RNA, purified RNA-protein complexes were resolved on 4-12% Bis-Tris gels and transferred to nitrocellulose membranes. Cells expressing only GFP were used as a negative control. The bottom panel shows a Western blot analysis of the samples probed with anti-GFP antibodies. **B:** Pie chart representing the distribution of Znf684-bound RNAs. **C:** Standardized metaplot profile showing the density of Znf684-iCLIP peaks, compared to the input RNA used as a control for background. CDS represents the coding sequence. **D:** Top two enriched sequence motifs found in Znf684 iCLIP-seq peaks. The E-values represent the significance of the motifs against randomly assorted control sequences. **E:** KEGG pathway enrichment analysis using genes that were targeted by Znf684. Only the top 10 most significant terms are shown (FDR ≤ 0.05). Darker nodes represent more significantly enriched gene sets. Bigger nodes represent larger gene sets. Thicker edges indicate more overlapping genes.

To identify which RNAs are bound by Znf684, we performed individual-nucleotide resolution UV crosslinking and immunoprecipitation (iCLIP) followed by high throughput sequencing (iCLIP-seq) experiments. iCLIP-seq peak distribution analysis indicated that the majority of peaks, ~57% (22,291/39,111 peaks), are located in the 3’UTRs of protein-coding genes (Supplementary Table S2; Figure 3B). ~35% of the total peaks were found in the coding regions of the target mRNAs, whereas only ~8% were found in the 5’UTRs (Figure 3B). Consistent with these observations, our metagene plots show that Znf684 binds to target mRNAs with a marked preference for 3’ UTRs (Figure 3C). We then performed motif analysis and found that the two most enriched motifs are guanosine (G)-rich sequences, both of which are very similar at the last three positions (Figure 3D). Collectively, these results suggest that Znf684 preferentially binds to the 3’UTRs of its target mRNAs, with a preference for G-rich sequences.

We next compared the iCLIP peak distribution with Znf684 chromatin occupancy. By utilizing our previously published chromatin immunoprecipitation combined with high throughput sequencing (ChIP-seq) data (13, 21), we identified 266 significant peaks for Znf684. These peaks were found across repeat elements (Supplementary Table S3) and did not overlap with iCLIP-seq peaks. Although Znf684 exhibits distinct DNA- and RNA-binding profiles, G-rich sequences appeared to be the preferred sites for binding in each case (Supplementary Figure S3).

To examine if Znf684 targets mRNAs with a specific function, we performed enrichment analysis using the Kyoto Encyclopedia of Genes and Genomes (KEGG). We found that Znf684 target mRNAs were significantly enriched in genes related to cellular macromolecule biosynthetic processes and organelle organization (FDR ≤ 0.05) (Figure 3E). Furthermore, regulation of gene expression and RNA processing were also over-represented in Znf684 targets (Figure 3E). These results suggest that Znf684 binds to functionally important transcripts *in vivo.*

### Znf684 has an RNA-binding profile similar to those of mRNA export factors

To further examine the functional link of Znf684 with mRNA export factors, we analyzed publicly available iCLIP-seq data for Nxf1 and Alyref (2). Our analysis indicated that both factors had RNA-binding profiles similar to that of Znf684, as they displayed a preference for binding to 3’ UTRs (Supplementary Figure S4A). Although Nxf1 and Alyref have been reported to bind RNA without any sequence preference (24, 25), our motif analysis identified moderately enriched G-rich sequence motifs for these proteins, similar to that of Znf684 (Supplementary Figure S4B). These observations are consistent with the idea that Znf684 functions in conjunction with mRNA export factors, including Nxf1 and Alyref, to regulate the export of target mRNAs.

### Znf684 affects mRNA export

To further investigate Znf684’s localization in the cell and its role in mRNA export, we separated nuclear and cytosolic fractions using cells overexpressing either GFP-Znf684 or free GFP. We overexpressed Znf684 in HEK293 cells and confirmed approximately twofold overexpression by reverse transcriptase quantitative polymerase chain reaction (RT-qPCR) (Figure 4A). Importantly, the expression levels of Nxf1 and Alyref remained unchanged upon Znf684 overexpression (Figure 4A). To rule out the possibility of cross-contamination between nuclear and cytosolic compartments, Western blotting experiments were performed, and the isolated nuclear and cytoplasmic fractions were probed with the nuclear U170k and cytoplasmic GAPDH markers. As shown in Figure 4B, we did not detect any cross-contamination between the cellular fractions (also see Supplementary Figure S5A). Furthermore, we also probed the Western blot with an anti-GFP antibody and found that Znf684 was present in both the cytoplasmic and nuclear fractions, consistent with our findings based on IF experiments (Figure 4B).

**Figure 4:**
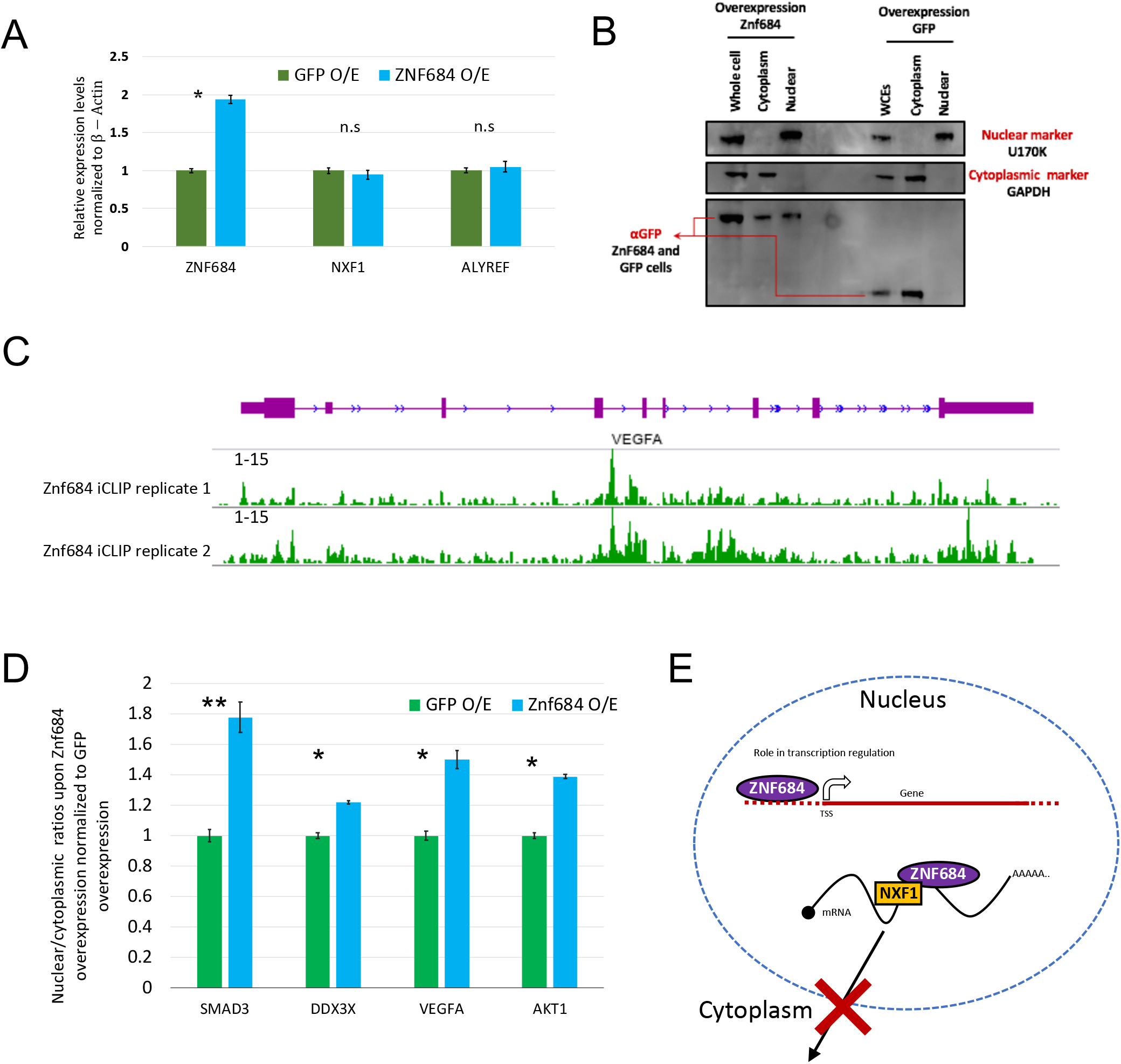
Znf684 affects mRNA export. **A:** Expression levels of Znf684, Nxf1 and Alyref mRNA in HEK293 cells expressing either free GFP or GFP-Znf684, as assessed by qRT-PCR. The qRT-PCR data were normalized to beta-ACTIN. * indicates t-test P ≤ 0.05. The individual error bars indicate the standard deviation for each sample. **B:** Western blotting analyses using nuclear and cytosolic fractions prepared from HEK293 cells expressing either free GFP or GFP-Znf684. The blots were probed with the indicated antibodies, with U170k acting as a nuclear marker and GAPDH acting as a cytoplasmic marker (also see Supplementary Figure S5 for replicate 2). **C:** Genome browser view for VEGFA gene showing the Znf684 iCLIP signal. **D:** RT-qPCR analysis to examine the abundance of target transcripts in the nucleus relative to cytoplasm upon GFP or Znf684-GFP overexpression. Statistical testing was performed utilizing Student’s t test: **p < 0.01. RT-qPCR experiments were performed in biological triplicate, and normalization was conducted relative to beta-Actin. The individual error bars indicate the standard deviation for each sample. **E:** Proposed models for Znf684 functions in the mRNA nuclear export pathway and transcription regulation. The nucleus is represented by a dotted circle, a target gene is represented in red, a target mRNA is represented by a curved black line, and the export factors are represented as coloured ovals and rectangles. The crossed-out arrow represents the downregulation of nuclear export.

Next, we compared the relative mRNA levels of four Znf684-targets, as identified by the iCLIP-seq data, in the nuclear and cytoplasmic fractions from GFP-Znf684 and GFP overexpressing cells by RT-qCPR experiments (Figure 4C; Supplementary Figure S5B). Our results show that the relative abundance of SMAD3, VEGFA, DDX3X and AKT1 transcripts in the nucleus was significantly (p<0.05) increased in the Znf684-overexpressing cells compared to the controls (Figure 4D). This indicates that fewer SMAD3, VEGFA, DDX3X and AKT1 mRNA transcripts reach the cytoplasm when Znf684 is overexpressed. These results indicate that Znf684 may play an inhibitory role in mRNA nuclear export (Figure 4E). The observed increase in the nuclear abundance of certain transcripts could also be related to transcription upregulation and/or mRNA degradation in the cytoplasm. Since Znf684 belongs to the KRAB family of C2H2-ZnF transcription factors, we investigated whether the expression levels of target genes were affected in cells overexpressing GFP-Znf684. Our results indicate that mRNA transcript levels decreased significantly (* p<0.05, ** p<0.01) upon Znf684 overexpression for AKT1, FHIT and VEGFA and SMAD3 (Supplementary Figure S5C). Thus, the observed increase in the nuclear transcript levels in Znf684-overexpressing cells is unlikely due to transcription upregulation. Further studies—including cellular fractionation followed by RNA sequencing and mRNA stability analyses—are currently under way to investigate the genome-wide effects of Znf684 overexpression on mRNA export.

### Reduced Znf684 expression in human cancers

Next, we examined potential links between Znf684 and cancer. Using archived patient RNA-seq data from ‘The Cancer Genome Atlas’ (TCGA), we examined the relationship of Znf684 expression levels to cancer diagnosis and prognosis (26). Our analysis revealed that Znf684 expression is significantly altered in 9 of the 31 cancer types we analyzed. Only three cancer types showed elevated Znf684 expression in tumors compared to normal tissue, namely Glioblastoma multiforme (GBM), Lymphoid Neoplasm Diffuse Large B-cell Lymphoma (DLBC), and Thymoma (THYM) (Supplementary Figure S6). We also found that Znf684 expression is significantly reduced in 6 of the 31 cancer types, including Kidney renal clear cell carcinoma (KIRC), Pheochromocytoma and Paraganglioma (PCPG), Prostate adenocarcinoma (PRAD), Thyroid carcinoma (THCA), Uterine Corpus Endometrial Carcinoma (UCEC), and Uterine Carcinosarcoma (UCS) (Figure 5A). These results suggest that altered expression of Znf684 might be a pathogenic event in certain cancer types.

**Figure 5:**
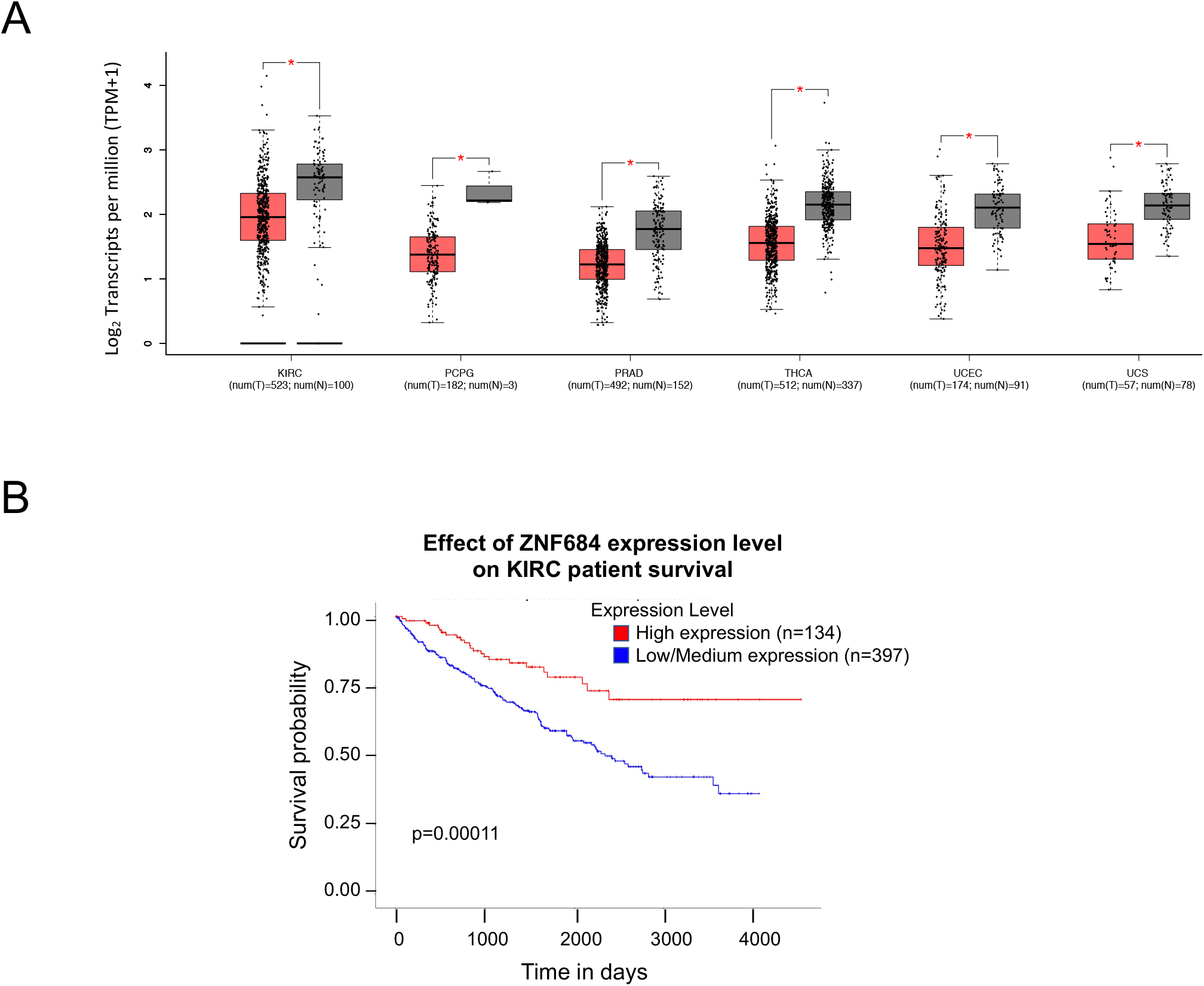
Znf684 expression is altered in various cancer types. **A:** Expression levels of Znf684 in normal and tumour cell samples. The plots are based on patient RNA-seq data available through TCGA. Box plots are shown for those cancer types for which Znf684 expression was significantly reduced in comparison to the normal tissues (also see Supplementary Figure S6). P-value cut off < 0.01. Note: The plots were generated using GEPIA (http://gepia.cancer-pku.cn/). **B**: Kaplan– Meier curve of overall survival in patients with KIRC using TCGA data. Patients were grouped according to the median expression of Znf684 using log-rank tests.

Further investigation using Kaplan–Meier curves (26) revealed that patients that have low expression of Znf684 exhibited significantly worse overall survival among those who were diagnosed with KIRC (P < 0.001) (Figure 5B). These results suggest that lower than normal expression level of Znf684 may be a predictor for worse overall survival for patients with KIRC. We suggest that continuing to study Znf684 might yield valuable insights for the development of cancer diagnostics or anti-cancer therapeutics.

## Discussion

mRNA export regulation plays an important role in the modulation of eukaryotic gene expression (7). In this study, we found evidence that Znf684 functions in mRNA nuclear export through a physical interaction with Nxf1. This idea is supported by its interactions with other nuclear export factors, similarities in RNA-binding profiles with those factors, and effect on mRNA export into the cytoplasm when overexpressed.

The NXF 1:NXT1 complex is a major export factor for bulk mRNAs (7). Additional factors, such as SR proteins, have also been shown to participate in mRNA export (27). Increasingly, it is becoming apparent that the nuclear export of specific classes of mRNAs can be selective (5, 28–30). In this context, we have recently shown that MKRN2, an RNA-binding E3 ubiquitin ligase, specifically binds to mRNAs that contain CU-rich regions in the 3’ UTR and regulates their export (31). Results presented in the current study point toward a role for Znf684 as an adapter for Nxf1 to regulate the export of specific mRNAs. Although many of the mechanistic details remain elusive, our observation that Znf684 overexpression inhibits export from the nucleus of certain mRNA transcripts suggests that it may have an inhibitory role in the nuclear export of a subset of mRNAs. Further experiments are necessary to explicitly elucidate detailed aspects of the mechanism by which Znf684 influences mRNA export. It also remains to be seen whether the DNA- or RNA-binding ability of Znf684 has a role in the observed expression changes. To this end, correlational analyses of iCLIP-seq, ChIP-seq, and knock-down RNA-seq data for Znf684 are under way.

Mitochondrial translation is another potential topic for future research on Znf684, as our AP-MS data showed that Znf684 interacts with many mitochondrial ribosomal proteins. Consistently, our IF experiments using Mitotracker showed that Znf684 localizes to the mitochondria. In line with Znf684 having a function in mitochondrial biology, the KEGG pathway analysis of our iCLIP data indicated that Znf684 binds to a subset of mRNAs that are involved in organelle organization. Together, these results point towards Znf684 not only having a role in the mRNA nuclear export pathway, but also having some function in the mitochondria. This function could be related to mitochondrial gene translation, given the association with mitochondrial ribosomal proteins.

The role of transcription factors in cancer is being increasingly recognised. Several C2H2-Znf family proteins, including Sp1, Znf281 and Zbtb7a, have been implicated in carcinogenesis (32–35). Reduced expression of Znf684 in several cancers and its correlation with poor patient survival in KIRC suggests that expression of Znf684 might have a negative impact on cancer cell proliferation. The potential role of Znf684 in cancer could be related to its function as a transcription factor, an RNA-binding nuclear export protein, or perhaps both. Further studies are required to dissect these possibilities.

## Experimental Procedures

### Cell cultures

HEK293 cells (Flp-In 293 T-REx cell lines) were obtained from Life Technologies (R780-07) (Invitrogen). Cell cultures were maintained in Dulbecco’s modified Eagle’s medium (DMEM) (Wisent Bioproducts catalogue number 319-005-CL), which was supplemented with 10% FBS (Wisent Bioproducts catalogue number 080-705), sodium pyruvate, non-essential amino acids, and penicillin/streptomycin as described (36). Doxycycline (1 μg/ml) was added to the culture medium 24 h before harvesting to induce the expression of the GFP-tagged proteins of interest.

### Antibodies

The following antibodies were used in this work: GAPDH (Abcam ab8245), U170k (Abcam ab83306), GFP (Life Technologies G10362), FLAG (Sigma monoclonal antibody catalogue number F1804), PABPC1 (Abcam ab21060), NXF1 (Abcam ab129160), and DEK (Abcam ab221545).

### Co-immunoprecipitations (Co-IPs)

Co-IPs were performed as described previously (37, 38). Cell pellets were lysed in lysis buffer (140mM NaCl, 10mM Tris pH 7.6–8.0, 1% Triton X-100, 0.1% sodium deoxy-cholate, 1mM EDTA) containing protease inhibitors. Cell extracts were incubated with 75 units of Benzonase (Sigma E1014) for 30 minutes in the cold room with end-to-end rotation. Cell debris was separated by microcentrifuging at 15,000g for 30 minutes at 4°C. The supernatant was incubated with 2.5μg of GFP antibody (Life Technologies G10362) overnight, and then 25μL protein G beads were added and incubated for an additional 2 hours with end-to-end rotation at 4°C. The beads were separated using a magnetic rack, and then the supernatant was removed. Next, the beads were resuspended in 1mL lysis buffer containing an additional 2% NP40 and 1% Triton X and rotated for 5 minutes in a cold room. This step was repeated three times. The samples were then boiled in SDS gel sample buffer. Samples were resolved using 10% SDS-PAGE and transferred to a PVDF membrane using a Gel Transfer Cell. Primary antibodies were used at 1:5000 dilution, and secondary antibodies were used at 1:10,000. Blots were developed using ECL Western Blotting Substrate.

### Affinity purification and Mass Spectrometry (AP-MS)

The AP-MS procedure was performed essentially as described (21, 39). Briefly, two independent batches of ~20×10^6^ cells were grown representing two biological replicates. Expression of the tagged protein was induced using doxycycline 24 h prior to harvesting. HEK293 cell pellets were lysed in high-salt NP-40 lysis buffer (10 mM Tris-HCl pH 8.0, 420 mM NaCl, 0.1% NP-40, plus protease/phosphatase inhibitors) with three freeze-thaw cycles. The lysate was sonicated and treated with Benzonase for 30 min at 4°C with end-to-end rotation. The cell lysate was centrifuged to pellet any cellular debris. We immunoprecipitated GFP-tagged Znf684 with 1 μg anti-GFP antibody (G10362, Life Technologies) overnight followed by a 2-hour incubation with Protein G Dynabeads (Invitrogen). The beads were washed 3 times with buffer (10mM TRIS-HCl, pH7.9, 420mM NaCl, 0.1% NP-40) and twice with buffer without detergent (10mM TRIS-HCl, pH7.9, 420mM NaCl). The immunoprecipitated proteins were eluted with NH4OH and lyophilized. Proteins for MS analysis were prepared by in-solution trypsin digestion. Briefly, protein pellets were resuspended in 44uL of 50mM NH4HCO3, reduced with 100mM TCEP-HCL, alkylated with 500mM iodoacetamide, and digested with 1 μg of trypsin overnight at 37°C. Samples were desalted using ZipTip Pipette tips (EMD Millipore) via standard procedures. The desalted samples were analyzed with an LTQ-Orbitrap Velos mass spectrometer (ThermoFisher Scientific). Raw MS data were searched with Maxquant (v.1.6.6.0) (40), yielding spectral counts and MS intensities for each identified human protein. The resulting data were filtered using SAINTexpress to obtain confidence values utilizing our two biological replicates. The AP-MS data generated from HEK293 cells expressing GFP alone were used as negative control for SAINT analysis. Data were represented as network using Cytoscape (V3.4.0) (41). Individual nodes were managed so as to arrange individual proteins into protein complexes.

### iCLIP-seq experiments

Individual nucleotide resolution UV crosslinking and immunoprecipitation (iCLIP) was performed as previously described (42) with the modifications detailed in our previous report (37). Briefly, cells were grown in 15cm culture plates and were UV cross-linked with 0.4J/cm2 at 254nm in a Stratalinker 1800 after induction with Doxycycline (1μg/ml) for 24 hours. Cells were lysed in 2mL of iCLIP lysis buffer. 1mL of lysate was incubated with 4 μL Turbo DNase (Life Technologies catalogue number AM2238) and 20 μL RNase I (1:250; Ambion catalogue number AM2294) for 5 minutes at 37°C to digest the genomic DNA and obtain RNA fragments within an optimal size range. GFP-Znf684 was immunoprecipitated using 5 μg of anti-GFP antibody (Life Technologies G10362). A total of 2% input material was obtained for size-matched control libraries (SMI) prior to the IPs. Following stringent washes with iCLIP high salt buffer and dephosphorylation with T4 polynucleotide kinase, on-bead-ligation of pre-adenylated adaptors to the 3’-ends of RNAs was performed using the enhanced CLIP ligation method (43). The immunoprecipitated RNA was 5’-end-labeled with 32P using T4 polynucleotide kinase (New England Biolabs catalogue number M0201L), separated using 4–12% BisTris-PAGE, and transferred to a nitrocellulose membrane (Protran). For the input sample, the membrane was cut to match the size of the IP material. RNA was recovered by digesting proteins using proteinase K (Thermo Fisher catalogue number 25530049) and subsequently reverse transcribed into cDNA. The cDNA was size-selected (low: 70 to 85 nt, middle: 85 to 110 nt, and high: 110 to 180 nt), circularized to add an adaptor to the 5’-end, linearized, and then PCR amplified using AccuPrime SuperMix I (Thermo Fisher catalog number 12344040). The final PCR libraries were purified on PCR purification columns (QIAGEN), and the eluted DNA was mixed at a ratio of 1:5:5 from the low, middle, and high fractions and submitted for sequencing on an Illumina HiSeq2500 to generate single-end 51 nucleotide reads with 40M read depth.

### Cellular Fractionation

Fractionation was performed as described (31). Cell pellets were resuspended in 1X PBS, then 10% of the cells were separated and pelleted by microcentrifuging at 400g. The remaining 90% was kept as the total fraction. The 10% cells were resuspended in 1X φ buffer (150mM potassium acetate, 5mM magnesium acetate, and 20mM HEPES pH 7.4), and 1X φ-plus buffer (1X φ buffer with the following added before use: 1mM DTT, 0.01% sodium deoxycholate, 0.25% Triton X, and protease inhibitor cocktail (Roche catalogue number 05892791001)). The mixture was iced for 3 minutes, and microcentrifuged for 5 minutes at 400g. The resulting supernatant served as the cytoplasmic fraction. The nuclear and endoplasmic reticulum (ER) fractions remained in the pellet, which was resuspended with 1X φ buffer and 1X φ plus buffer, then microcentrifuged for 5 minutes at 400g. The supernatant containing the ER was removed, and the pellet was washed with 1X φ buffer. The resulting pellet containing the nuclear fraction was resuspended in 1X φ buffer. Samples were resolved using 10% SDS PAGE and probed with nuclear (U170K, Abcam ab83306) marker antibodies at 1:1000 dilution and cytoplasmic (GAPDH, Abcam ab8245) marker antibodies at 1:10,000 dilution.

### Immunofluorescence microscopy (IF)

IFs were performed as described (37). GFP-Znf684 and GFP expressing HEK293 cells were seeded on poly-L-lysine coated and acid-washed coverslips. Protein expression was induced using 1μg/ml doxycycline for 24 hours. Cells were washed three times with PBS, then fixed in 4% Paraformaldeyde for 15 minutes. Cells were subsequently permeabilized with 0.2% Triton X-100 in PBS for 5 minutes and incubated with block solution (1% goat serum, 1% BSA, 0.5% Tween-20 in PBS) for 1 hour. GFP antibody was used for staining at 1:100 concentration in block solution for 2 hours at room temperature (RT). Cells were incubated with Goat anti-mouse secondary antibody and Hoescht stain in block solution for 1 hour at room temperature. Cells were fixed in Dako Fluorescence Mounting Medium (S3023). Imaging was performed the next day using a Zeiss confocal spinning disc AxioObserverZ1 microscope equipped with an Axiocam 506 camera using Zen software. A single focal plane was imaged.

### Reverse Transcription Quantitative Polymerase Chain Reaction (RT-qPCR)

RT-qPCR was performed as described (31). Total RNA was extracted using Trizol Reagent (Invitrogen catalogue number 15596018) as per the manufacturer’s instructions. RNA was used for cDNA synthesis with the SuperScript VILO Kit (Invitrogen catalogue number 11754). RT-qPCR was performed with Fast SYBR Green Master qPCR mix (Applied Bioscience 4385617) on an Applied Biosystems 7300 real time PCR System (Thermo Fisher catalogue number 4406984). For qPCR, the following program was used: 40 cycles of 95°C for 15 s and 60°C for 30 s, with a final cycle of 95 °C for 15 s and then 60 °C. Actin was used as loading control. The data was analyzed using the ΔΔCT method as described previously (44). Primer sequences are provided in Supplementary Table S2.

### TCGA data analysis: Differential expression analysis and Kaplan–Meier plots

Gene expression analysis was performed using the online tool ‘UALCAN’ (http://ualcan.path.uab.edu/index.html) (14) and GEPIA (45). These tools give access to a large collection of data including ‘The Cancer Genome Atlas (TCGA)’. Differential expression and survival analyses of Znf684 in tumor/normal tissue from renal clear cell carcinoma (KIRC) were performed.

### Publicly available datasets

Publicly available datasets were obtained for IF images and RNA-seq expression values from the Human Protein Atlas (proteinatlas.org, last accessed April 8, 2021) and VastDB (vastdb.crg.eu, last accessed April 8, 2021), respectively. iCLIP data for mRNA export factors were acquired from GSE113896 (2). ChIP-seq and AP-MS data were acquired from Gene Expression Omnibus (GEO; http://www.ncbi.nlm.nih.gov/geo/) under accession number GSE76496 and PRIDE (https://www.ebi.ac.uk/pride/archive/) under accession number PXD003431, respectively.

## Data availability

CLIP-seq data generated can be found online at Gene Expression Omnibus (GEO, https://www.ncbi.nlm.nih.gov/geo/).

## Acknowledgements

We would like to thank Nujhat Ahmed, Giovanni Livingston Burke, and Kristie Ng for their experimental help and advice. Ernest Radovani is gratefully acknowledged for helpful discussions and assistance with MS analyses. Guoqing Zhong is acknowledged for her support in tissue culture. We would also like to thank Edyta Marcon for her help in obtaining materials and administrative support.

## Author contributions

S.N-S. and J.F.G. conceived the project. A.N. performed co-IP experiments, RT-qPCR, participated in IF experiments, analyzed data and drafted the manuscript. S.N-S performed iCLIP-seq experiments, Data analysis, edited the manuscript and co-ordinated the project. S.P. analyzed iCLIP data. S.F. performed IF analysis. B.J.B. supervised SF and edited the manuscript. J.F.G. supervised the project, participated in data analysis, and edited the manuscript. This work was supported by Canadian Institutes of Health Research Foundation Grant FDN-154338 to JFG.

