## Supplemental Figure legends for "KRAB Zinc Finger protein Znf684 interacts with Nxf1 to regulate mRNA export"

**Supplementary Figure S1: Znf684 has low expression across cell lines.**

**A:** Analysis of relative expression of Znf684 across cell lines, measured in corrected Reads Per Kilobasepair and Million mapped reads (cRPKM). **B:** Relative expression of Znf684 across embryonic developmental stages. Note: The expression data was acquired from the Vastdb database (https://vastdb.crg.eu/gene/ENSG00000117010@hg38).

**Supplementary Figure S2: Znf684 localizes to both the nucleus and cytoplasm.**

IF analysis of GFP-Znf684 in various cell lines, including RH-30, SiHa, and U2-OS. IF data was acquired from the Human Protein Atlas (https://www.proteinatlas.org/).

**Supplementary Figure S3: Znf684 DNA-binding motif.**

The motif is based on ChIP-seq data published previously and is publicly available (http://kznfmotifs.ccbr.utoronto.ca/krab.php).

**Supplementary Figure S4: Nxf1 and Alyref have RNA-binding profiles similar to that of Znf684.**

**A:** Metagene analyses of Nxf1 and Alyref iCLIP-seq data from transcription start sites (TSS) to transcript end sites (TES) indicate enrichment near the 3'UTRs of target genes. Alyref is also enriched around the TSS. **B:** Top two enriched sequence motifs found in each of Nxf1 and Alyref CLIP-seq peaks. The E-values represent the significance of the motifs against control sequences.

**Supplementary Figure S5: Subcellular fractionation of cells overexpressing GFP or GFP-Znf684.** Western blotting analyses using nuclear and cytosolic fractions prepared from HEK293 cells expressing either free GFP or GFP-Znf684. The blots were probed with the indicated antibodies, with DEK acting as a nuclear marker and GAPDH acting as a cytoplasmic marker. **B:** Genome browser view for SMAD3 and DDX3X genes showing the iCLIP signal of Znf784. **C:** Expression levels of AKT1, FHIT, VEGFA and SMAD3. mRNA levels in HEK293 cells expressing either free GFP or GFP-Znf684 were quantified using qRT-PCR. The qRT-PCR data were normalized to beta-Actin expression. * indicates t-test $P \leq 0.05$, ** indicates $P \leq 0.01$. The individual error bars indicate the standard deviation for each sample.

**Supplementary Figure S6: Expression levels of Znf684 in normal and tumour cell samples.** The plots are based on patient RNA-seq data available through TCGA. Box plots are shown for those cancer types for which Znf684 expression was significantly increased in comparison to the normal tissues. P-value cut off $< 0.01$. The plots were generated using GEPIA (http://gepia.cancer-pku.cn/).
