## Supplemental figures for "KRAB Zinc Finger protein Znf684 interacts with Nxf1 to regulate mRNA export"

### Supplementary Figure S1

A

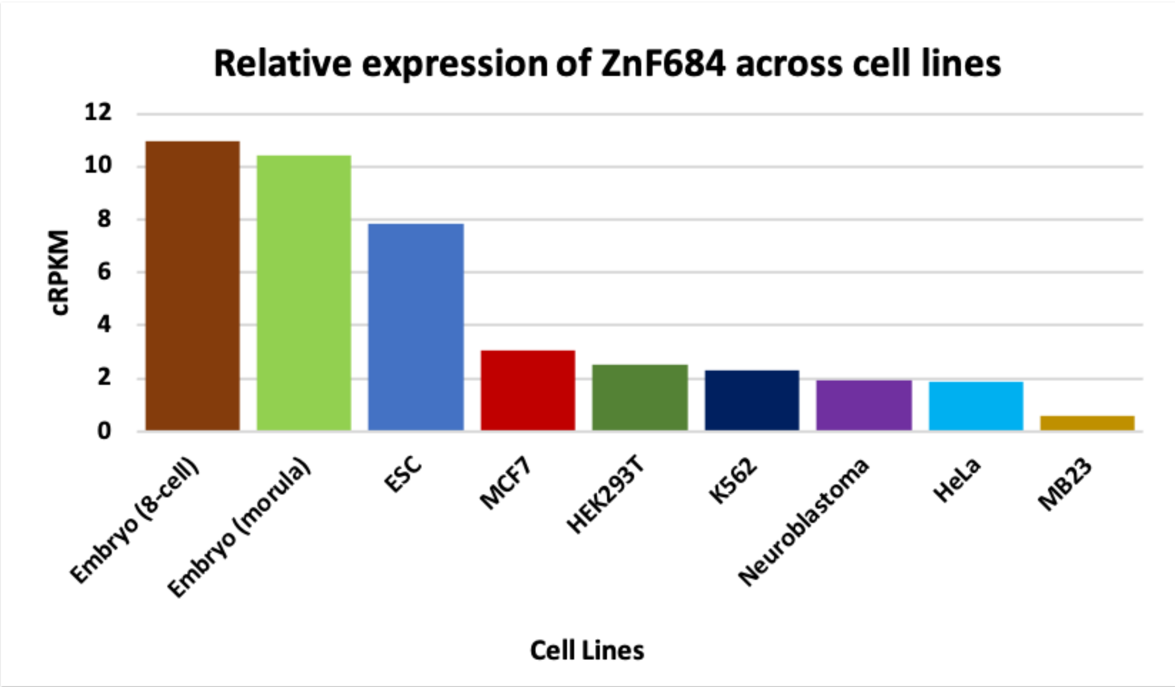


B

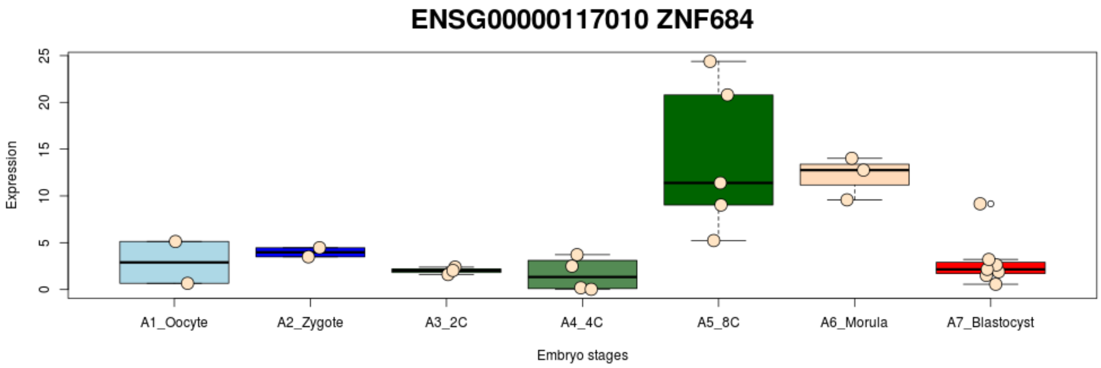

### Supplementary Figure S2

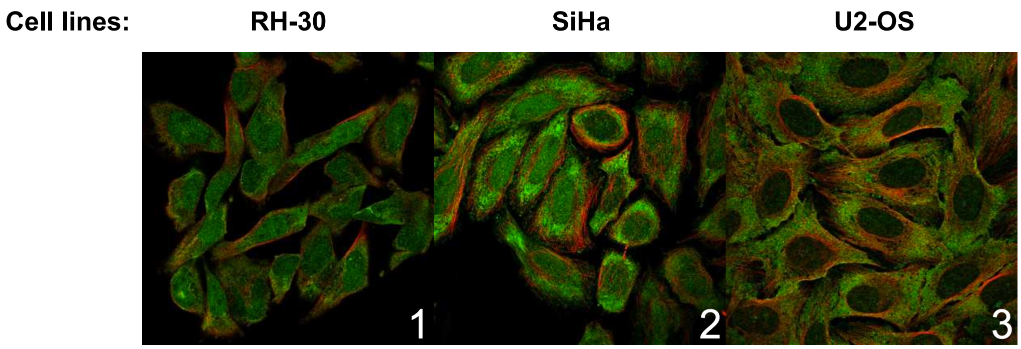

Supplementary Figure S3

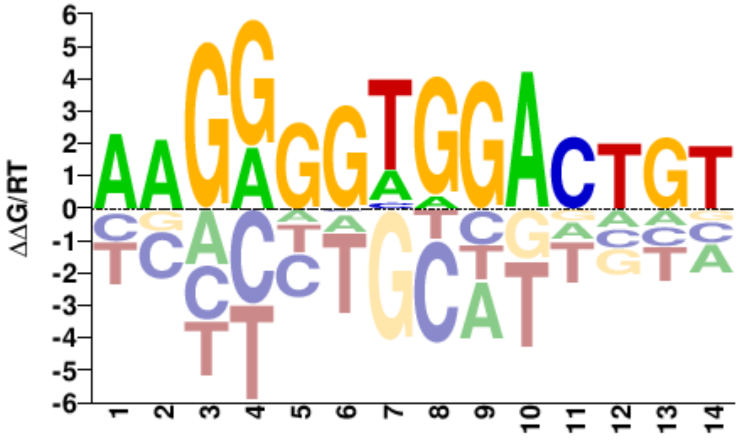

### Supplementary Figure S4

A

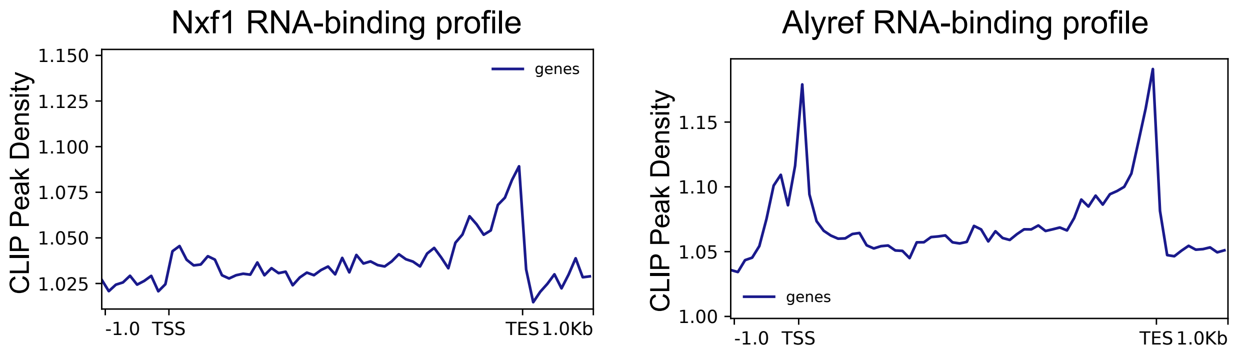


B

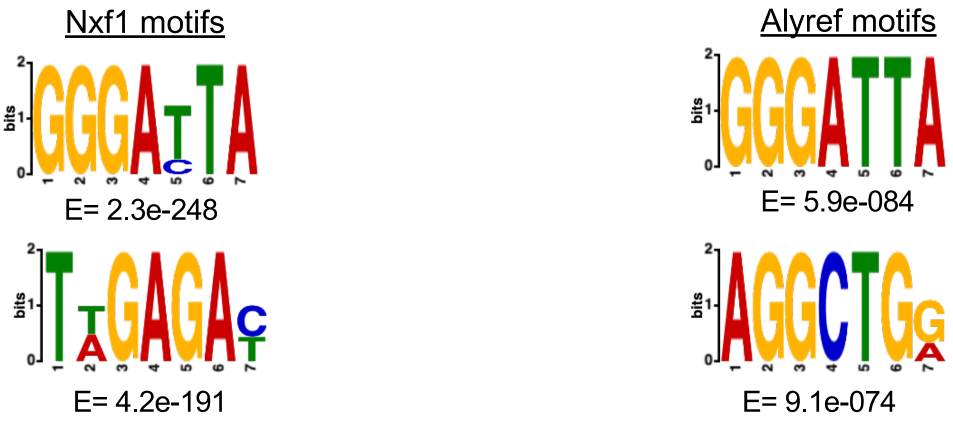

### Supplementary Figure S5

A

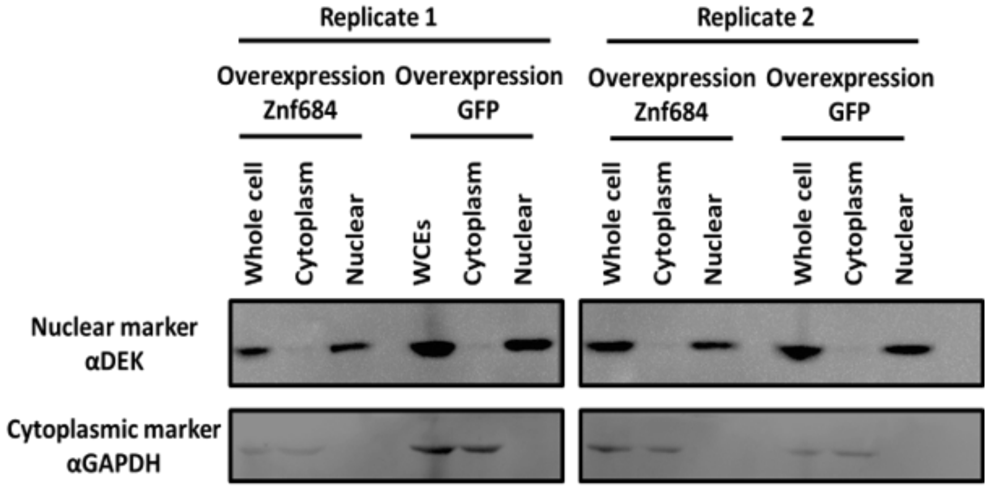


B

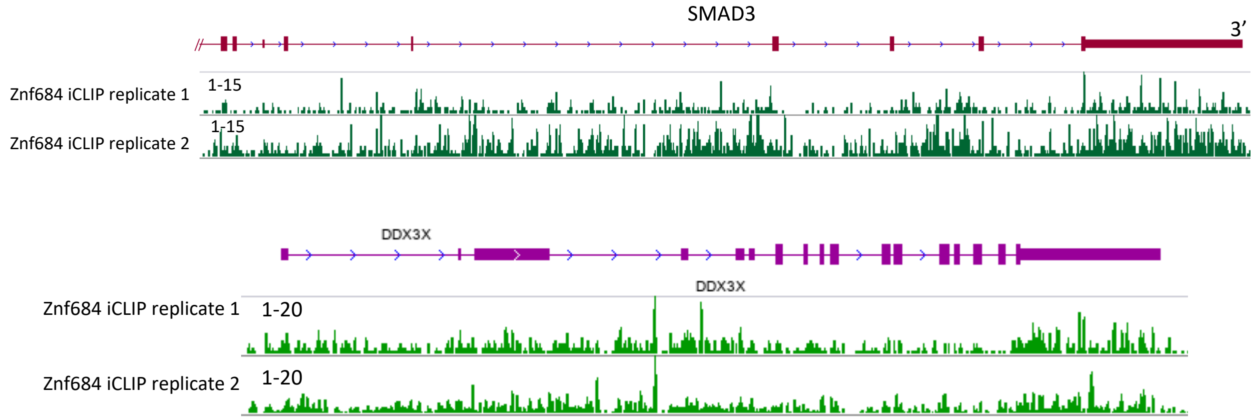


C

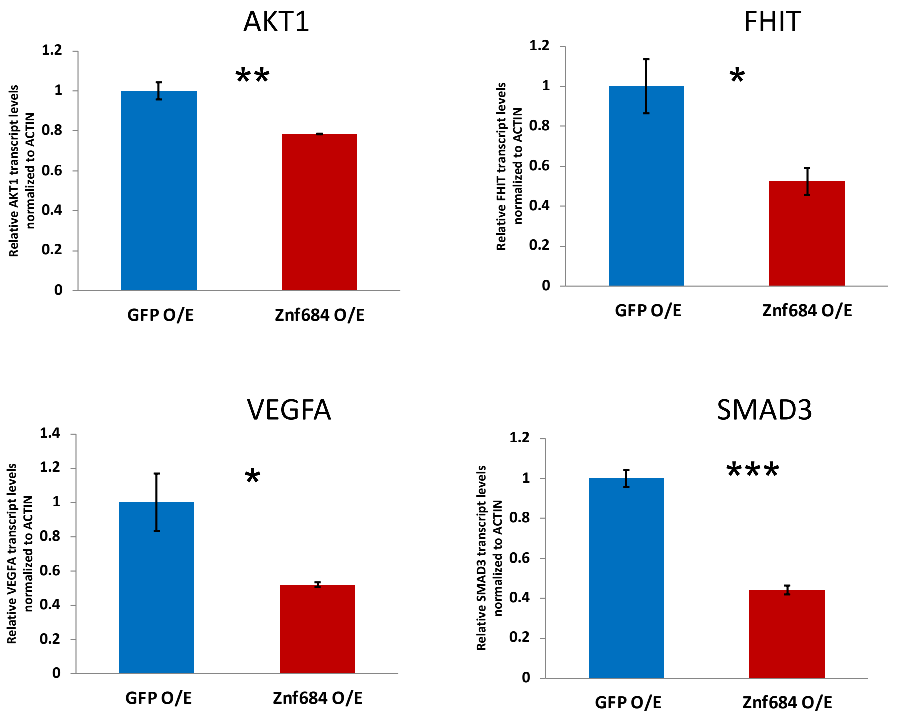

### Supplementary Figure S6

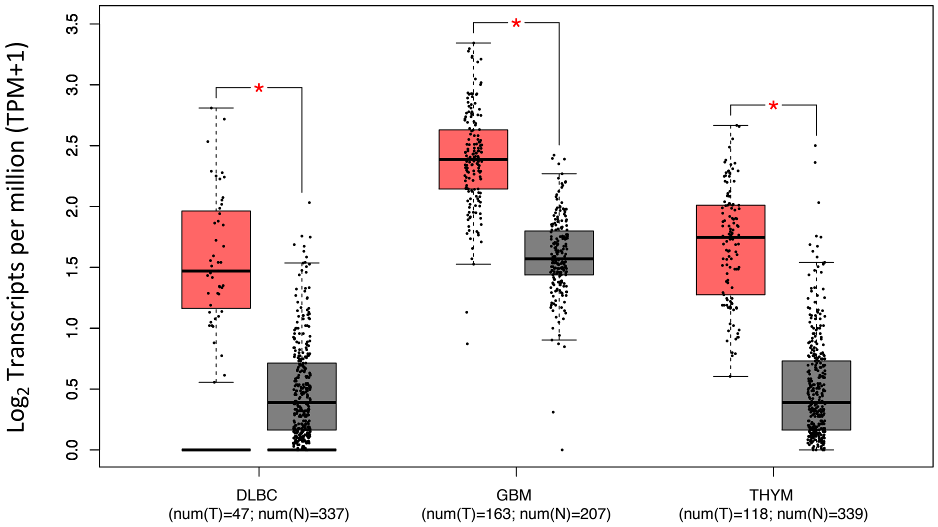
